# Phasing genome assemblies of non-model animal species in the era of high-accuracy long reads

**DOI:** 10.1101/2024.06.16.599187

**Authors:** Nadège Guiglielmoni, Philipp H. Schiffer

## Abstract

The technological shift brought by high-accuracy long reads is transforming the genomic landscape and bringing forth chromosome-scale reference assemblies across the tree of life. Large-scale initiatives have emerged to encourage the production of such reference genomes, with the Earth BioGenome Project (EBP) establishing rigorous quality standards, while consortia like the Vertebrate Genome Project (VGP), Darwin Tree of Life (DToL), and European Reference Genome Atlas (ERGA) focus on large-scale sequencing and assembly production.

By precisely discriminating alternative alleles and spanning multiple heterozygous regions, high-accuracy long reads enable assembly phasing, a more accurate representation of genomes than traditional collapsed assemblies. However, phasing assemblies remains challenging when access to long-read sequencing is restricted by low DNA material, or limited access to high-end sequencing facilities.

To address these limitations, we evaluate three distinct long-read strategies: the established PacBio HiFi standard, the portable and more accessible Nanopore R10.4.1 technology, and the PacBio HiFi ultra-low input protocol designed for samples with only nanograms of DNA. We benchmarked five assemblers using empirical reads of these three technologies for the genome of the parthenogenetic nematode *Plectus sambesii*. Through a comprehensive assessment of contiguity, structural correctness, and gene and transposable element content, we identified substantial differences in haplotype separation, further validated across five additional non-model animal species for which reads from one sequencing technology were available. This study provides critical guidelines for *de novo* phased assembly. Ultimately, we show that the feasibility of haplotype-resolved assembly does not lie in the choice of sequencing technology, but in the selection of the appropriate assembler.

## Introduction

High-accuracy long reads have brought drastic improvements to the field of genomics, raising the bar for reference assemblies in terms of quality and contiguity. Pacific Biosciences (PacBio) introduced the first long reads with a random error pattern below 1% through Circular Consensus Sequencing. This technology is a development of the previous PacBio protocol to pass over a fragment multiple times and produce high-fidelity reads, then designated as PacBio HiFi reads [1], while maintaining a length around 15 kilobases (kb). In parallel, Oxford Nanopore introduced the R10.4.1 chemistry to mitigate the issue of non-random homopolymeric errors [2]. The resulting reduction of error rate rendered Nanopore R10.4.1 reads sufficient for *de novo* assembly without the need of additional high-accuracy data for polishing [3]. These reads now routinely reach Q20+ accuracy and can span hundreds of kilobases [4].

PacBio HiFi and Nanopore R10.4.1 reads are opening new possibilities for haplotype phasing. Their low error rate allows for the discrimination of alternative alleles, and their length can span multiple consecutive heterozygous regions, facilitating correct allele association within contigs. Long-read assemblies are usually scaffolded using genome-wide chromosome conformation capture (Hi-C), which uses the density of 3D contacts to order and orient contigs [5]. Hi-C scaffolding further assists in resolving haplotypes by distinguishing high-frequency *cis* interactions (within the same chromosome) from lower-frequency *trans* interactions (between homologous chromosomes) [6]. With recent progress in separating alleles using high-accuracy long reads, alternative haplotypes are now commonly released along collapsed assemblies in public repositories such as the European Nucleotide Archive [7]. The Vertebrate Genome Project [8] established a complete assembly pipeline, suppported by the Galaxy platform [9], which includes the generation of phased assemblies [10]. This pipeline generally relies on PacBio HiFi and Hi-C reads and the assembler hifiasm [11], and has been largely applied for vertebrate genome assembly projects, which typically have a low heterozygosity. Conversely, the European Reference Genome Atlas [12] adopted a more heterogeneous approach, incorporating both Nanopore and PacBio HiFi to concurrently accommodate existing regional sequencing infrastructures, and the broader range of genomic complexities and heterozygosity levels found across diverse eukaryotic taxa.

Current guidelines from the Earth BioGenome Project [13] advocate for the use of either PacBio HiFi or Nanopore R10.4.1 sequencing as the foundation for initial genome assemblies. While major centralized consotia (i.e. the Vertebrate Genome Project, the Darwin Tree of Life [14]) predominantly rely on the PacBio HiFi platform, the large infrastructure required for such sequencing can be prohibitive for smaller research initiatives. This creates a significant technological divide, particularly in the Global South, where the logistical and financial burden - from the million dollar acquisition of a Revio sequencer, to dedicated supporting staff and computational resources — limits the feasibility of independent projects [15]. In contrast, the portability and lower entry costs of Nanopore devices offer a more democratized alternative, essential for empowering local initiatives to lead biodiversity genomics in their own regions [15, 16]. Consequently, it is crucial to assess whether both sequencing technologies can yield comparably high-quality phased assemblies, as a prerequisite not only for biological exactitude, but also for addressing societal stakes of scientific equity.

A major barrier to phased assemblies in biodiversity genomics lies in the DNA input requirement. Traditional protocols often rely on large amounts of fresh tissue for long-read sequencing and Hi-C, which is impractical for small or rare specimens [17]. For collapsed assembly, sequencing data from different individuals can be combined, although this can complicate the assembly process. For phased assembly, adding individuals would multiply the number of haplotypes to resolve. With such limitations, genome assembly projects may rely exclusively on long reads with no possibility to generate complementary long-range data, such as Hi-C, from a single individual. Furthermore, the requirements for long-read sequencing alone (over one microgram of DNA, or at least, some hundreds of nanograms) may not be achievable for small animals. To tackle this challenge, several amplification protocols have been released (PiMmS [18], AmpliFi [19]) to produce PacBio HiFi sequencing reads from only one nanogram of DNA.

In response to technological developments, novel genome assembly methods were developed to leverage the information provided by high-accuracy long reads. Some already existing low-accuracy long-read assemblers were updated to incorporate PacBio HiFi and Nanopore R10.4.1 reads, such as HiCanu [20], a newer version of Canu [21] implementing PacBio HiFi reads, and Flye [22] which added dedicated parameters for Nanopore R10.4.1 and PacBio HiFi reads. Other assemblers were developed specifically for high-accuracy long reads. hifiasm [11] implemented a string-graph approach to assemble PacBio HiFi reads and sequentially added Hi-C [23] and ultralong Nanopore [24] reads to improve contig resolution. Verkko [25] built upon the Canu assembler combined with Multiple Bipartite Graphs to also integrate ultra-long Nanopore and Hi-C reads with PacBio HiFi *de novo* assembly. Recently, hifiasm incorporated a novel read-phasing correction step suited for Nanopore R10.4.1 reads and their remaining systematic homopolymeric errors [3]. Alternatively, the assembler PECAT [26] proposed a comprehensive pipeline for PacBio HiFi and Nanopore reads with an overlap-based correction step and assembly yielding initally collapsed contigs. Alleles are reconstructed *post hoc* through variant calling and a second assembly step.

The field of genome assembly has now reached a turning point: when haplotype separation is feasible and yet still difficult, should every assembly project aim for a phased assembly when haplotype divergence is not relevant to the research question being addressed? Collapsed reference assemblies have been the goal so far for all species, including those with mid (>1%) to high (>3%) heterozygosity levels. But chosing one allele to represent each heterozygous site means dismissing information from alternative alleles and how these alleles associate. Inaccurate identification and purging of alternative haplotypes has previously led to suboptimal pseudohaploid assemblies where heterozygous regions were artificially duplicated [27]. Furthermore, collapsing haplotypes forces the juxtaposition of alleles from distinct homologous chromosomes, creating a mosaic chimeric sequence. This haplotype-switching interferes with Hi-C scaffolding, as a chimeric consensus inevitably yields conflicting contact signals and can induce artifacts in contact maps, subsequently requiring time-consuming manual curation [28]. This issue is particularly pronounced in genomes with high heterozygosity; therefore, phased assemblies are essential to provide the most accurate genomic representation, even when only a single haplotype is retained for downstream analyses.

In this study, we aimed to evaluate the potential of different long-read sequencing technologies and assemblers to yield high-quality phased assemblies for non-model animal species. First, we conducted a proof-of-principle comparison of sequencing technologies and assemblers on one parthenogenetic nematode species, *Plectus sambesii*, which has a small genome size and a high heterozygosity level. We generated non-amplified Nanopore and PacBio HiFi reads with a large input of DNA, and ultra-low-input (ULI) amplified PacBio HiFi reads, starting with less than a nanogram of DNA. These long reads were assembled using five programs, namely Canu [21, 20], Flye [22], hifiasm [11], PECAT [26] and Verkko [25]. Assemblies were evaluated for their size, contiguity, and structural correctness. We implemented a comprehensive methodology to assess phasing which repurposes state-of-the-art evaluation programs by collecting haplotype-relevant statistics. We assessed assembly size and contiguity, structural correctness, ortholog and *k*-mer completess, and representation of transposable elements. In a second step, we aimed to confirm whether the patterns observed in long-read assemblies of *Plectus sambesii* also applied to other non-model animal species. Assemblers were thus benchmarked on several species for which either Nanopore R10.4.1 or PacBio HiFi reads were publicly available. We indeed identified variability in phasing efficiency between sequencing technologies and assemblers, alike the patterns initially identified in *Plectus sambesii*. Our analysis provides an overview of strategies to yield haplotype-resolved *de novo* genome assemblies from highly accurate long reads and can serve as a guideline for genome assembly projects of non-model animal species.

## Material & Methods

### Long-read sequencing of *Plectus sambesii*

High-molecular-weight DNA was extracted from *Plectus sambesii* as described in [29]: from plates of cultured worms with a CTAB-phenol-chloroform protocol for Nanopore and non-amplified PacBio HiFi sequencing; and with a salting-out protocol on a single individual for ultra-low-input PacBio HiFi sequencing. The Nanopore library was prepared with the LSK114 kit and sequenced on two MinION flowcells. Reads were basecalled using Dorado v0.3.1 [30] in duplex mode with model dna_r10.4.1 e8.2_400bps_supv4.2.0 and the reads were converted to fastq using SAMtools v1.18 [31] with the module `samtools fastq`. The non-amplified PacBio HiFi libraries were prepared with shearing using the Megaruptor3, SPK3.0 library prep kit, and size-selection with AMPure PB beads or PippinHT. The libraries were sequenced on a Revio for 30h. The ultra-low-input amplified PacBio HiFi library was prepared by shearing the DNA in gTubes, using the SPK3.0 library prep kit with PCR combination of KOD + RM-B, and with size selection using AMPure PB beads. The library was sequenced on a Sequel IIe instrument with 30 hours movie time.

### Quality control

Quality and length of Nanopore reads and PacBio HiFi reads were plotted using NanoPlot v1.46.2 [32] on the Galaxy Europe platform [9] (usegalaxy.com). Genome size and heterozygosity were estimated using jellyfish v2.2.8 [33] and GenomeScope v2.0 [34] with *k* =21.

### *De novo* assembly

#### Plectus sambesii

Nanopore Q20 reads were selected using chopper v0.5.0 with parameter -q 20. The full Nanopore dataset and the Nanopore Q20 dataset were assembled separately using Canu v2.2 [21] with parameters `-nanopore -trimmed`, Flye v2.9.6 [22] with parameter `--nano-hq --keep-haplotypes`, hifiasm v0.25.0 [3] with parameters `-l 0 –ont` and PECAT v0.0.3 [26] with the ONT config file and parameter `genome size=250000000`. PacBio HiFi reads were assembled using Flye v2.9.6 [22] with parameter `--pacbio-hifi --keep-haplotypes`, hifiasm v0.25.0 [11] with parameter `-l 0`, and with and without Nanopore reads using the parameter `--ul` [24], and PECAT v0.0.3 [26] with the HiFi config file and parameter `genome size=250000000`. Verkko 1.3.1 [25] was run on Galaxy Europe [9], as we failed to install Verkko locally (including more recent versions), using PacBio HiFi reads (amplified or non-amplified) and with or without Nanopore reads. For PECAT assemblies, the primary and alternate assemblies were concatenated into a draft phased assembly subsequently labelled as “combined”.

#### Other species

Publicly available reads were collected from European Nucleotide Archive: *Erebia palarica* ERR12952298, *Xylophaga dorsalis* [35] ERR9744411, *Eunicella cavolini* ERR14694924, ERR14694925, ERR14694926, *Piscicola geometra* [36] ERR9744411, *Hypochthonius rufulus* [37] ERR15977476. Assemblers were run as described previously, with genome sizes set to: *Erebia palarica* 800 Mb, *Xylophaga dorsalis* 900 Mb, *Eunicella cavolini* 1000 Mb, *Piscicola geometra* 400 Mb, *Hypochthonius rufulus* 350 Mb.

### Decontamination

Bacterial contamination is expected for *Plectus sambesii* as its feeding source. BUSCO v5.8.0 [38] was run with parameters `-m genome -l nematoda odb12`, BLAST v2.10.0 [39] was run against the nt database with parameters `-outfmt “6 qseqid staxids bitscore std sscinames scomnames” -max hsps 1 -evalue 1e-25` and minimap2 v2.24 [40] using respective long reads with parameter `-ax map-ont` for Nanopore reads and `-ax map-hifi` for PacBio HiFi reads. The outputs were provided to BlobToolKit v4.3.2 [41] and contaminant bacterial and fungal contigs were removed. For hifiasm assemblies, contigs flagged as circular were also removed. Assemblies of *Xylophaga dorsalis* and *Piscicola geometra* were decontaminated in the same manner, except for the lineage provided to BUSCO v5.8.0 as `-l mollusca odb12 and -l metazoa odb12`, respectively. For *P. geometra*, only contigs flagged as Euglenozoa or Proteobacteria were removed, to avoid removing potential host contigs and affect downstream completeness evaluation. Other species were assessed but decontamination was not deemed necessary.

### Assembly evaluation

Basic statistics were computed for all assemblies using gfastats v1.3.6 [42]. BUSCO v5.8.0 [38] was run as described previously, with lineages: Nematoda odb12 for *Plectus sambesii*, Lepidoptera odb12 for *Erebia palarica*, Mollusca odb12 for *Xylophaga dorsalis*, Metazoa odb12 for *Eunicella cavolini* and *Piscicola geometra*, and Arachnida odb12 for *Hypochthonius rufulus*. KAT v2.4.2 [43] was run with the module `kat comp` and the Illumina dataset as reference for *Plectus sambesii*, or the long-read data used for assembly for other species. Proportions of 1X and 2X *k*-mers were collected from the log of `kat comp`. Repeats and transposable elements were identified using EDTA v2.0.1 [44] with parameters `--anno 1 –sensitive`. Long reads were mapped to the final assemblies using minimap2 v2.24 [40] as described previously; the output was sorted using SAMtools v1.18 [31] and provided as input to Sniffles2 v2.7.2 [45].

## Results

### Initial sequencing and assessment for *Plectus sambesii*

To first examine phasing efficiency with different sequencing and assembly methods on a small test genome, we chose to resequence and assemble the genome of the parthenogenetic nematode *Plectus sambesii*, which had previously been assembled from short Illumina reads [46]. This species has a small haploid genome size of 120 Mb with a heterozygosity of 3.80% (Supplementary Figure S1). As a diploid, the cumulative size of the two haplotypes is expected to reach 240 Mb. We generated non-amplified Nanopore R10.4.1 and PacBio HiFi reads, and amplified PacBio HiFi reads. Coverage depth per haplotype for each dataset reached: 46X for Nanopore R10.4.1 reads (N50: 35.1 kb) with 19X over Q20 (N50: 35.5 kb), 27X for non-amplified PacBio HiFi reads (N50: 12.8 kb), and 40X for amplified PacBio HiFi reads (N50: 15.8 kb). These sequencing depths match recommendations from the Earth Biogenome Projects, over 20X for Nanopore R10.4.1 and 12.5X for PacBio HiFi [47]. Quality and length analyses highlight the advantage of Nanopore reads in terms of length, while PacBio HiFi reads have higher quality values (Supplementary Figure S2).

### Submegabase-level contiguity achieved by high-accuracy long-read assemblies

To evaluate the quality of the assemblies, we first assessed their size and contiguity. The assembly size should be close to the expected genome size, that is, in the case of a phased assembly, the cumulative size of all haplotypes. Shorter assembly sizes may be owed to missing haplotypes and underrepresented repeats, and oversized assemblies can often be explained by *in silico* duplications and contamination. Contiguity is usually measured with the N50, which represents the largest fragment length in an assembly for which 50% of the assembly is represented in fragments of longer or equal length. This metric also has variations as N60, N70, N80 and N90 for 60%, 70%, 80% and 90%. The NG50 is similar to the N50, but is calculated against an expected genome size rather than the assembly size. Standard N50 metrics can be misleading in the context of diploid assembly, as a mostly collapsed assembly may appear contiguous while missing half of the genomic information. We therefore employed NG50 and NG90 for this analysis, which ensure that contiguity is measured relative to the total expected sequence length, thereby prioritizing haplotype-resolved contiguity.

All assemblies had a size close to the expected diploid genome size of 240 Mb (Figure 1), and they reached an NG50 over 1 Mb. Nanopore reads (all and Q20 only) resulted in assemblies with the highest contiguities, with NG50s of 19.2 and 14.5 Mb for hifiasm and PECAT (Nanopore) assemblies, an NG50 of 18.0 Mb for hifiasm (Nanopore Q20) and an oustanding NG90 of 16.7 Mb for hifiasm (Nanopore). All non-amplified PacBio HiFi assemblies had an NG50 over 6.8 Mb, and a highest NG50 of 18.0 Mb with hifiasm, close to the NG50s of the most contiguous Nanopore assemblies. The combination of Nanopore and non-amplified PacBio HiFi reads increased the NG50 of the Verkko assembly (from 6.8 to 18.0 Mb), and the NG90s of both hifiasm and Verkko assemblies (from 5.8 and 3.1 Mb, to 11.4 and 8.9 Mb). Amplified PacBio HiFi assemblies were overall less contiguous, with NG50s ranging from 1.1 to 1.7 Mb, and NG90s from 77 kb to 396 kb. The addition of Nanopore did improve contiguity of hifiasm and Verkko assemblies, but NG50s and NG90s remained lower than those of hifiasm and Verkko assemblies of non-amplified long reads.

**Figure 1:**
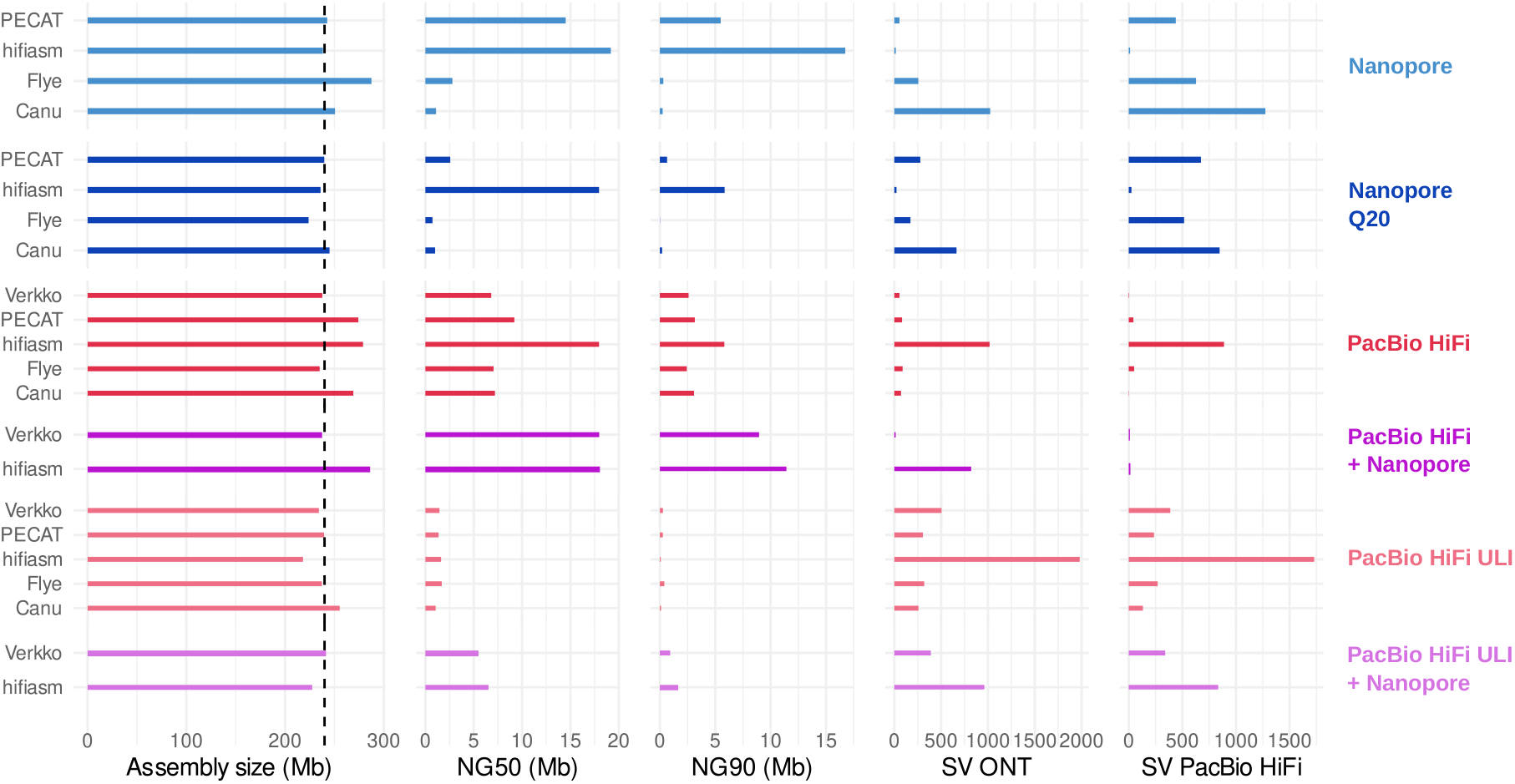
Basic statistics and structural correctness evaluation. Assembly size, NG50, and NG90 are presented from left to right. The NG50 and NG90 represent the assembly contiguity and were computed against an estimated size of 240 Mb (with both haplotypes included). Structural variants (SV) (right) were detected with the same Nanopore and non-amplified PacBio HiFi reads, as potential misassemblies.

### Variable structural correctness depending on sequencing technology and assembler combination

Achieving high contiguity is not sufficient to deem an assembly of high quality. In fact, the contiguity could be maximized by concatenating contigs, with no regard for the correctness of these junctions. To evaluate the structural correctness of the contigs, we searched for structural variants (SVs), i.e. insertions, deletions, translocations, inversions, duplications. These were detected using the same non-amplified Nanopore and PacBio HiFi reads which were used for assembly; hence we expected to find minimal amount of SVs, as they would constitute potential misassemblies. Variations in numbers of detected SVs between assemblies are usually similar, whether they were identified with Nanopore or PacBio HiFi reads. For Nanopore assemblies (Figure 1), Canu assemblies had the largest amount of SVs, with some decrease when only Nanopore Q20 reads were used (1024 SVs Nanopore, 1274 SVs HiFi for Nanopore assemblies; 664 SVs Nanopore, 847 SVs HiFi for Nanopore Q20 assemblies). hifiasm had the lowest quantity of SVs, up to a maximum of 25 detected SVs across detection and assembly datasets. By contrast, hifiasm led to the highest numbers of SVs for PacBio-HiFi-based assemblies, both for amplified and non-amplified data (1980 SVs Nanopore, 1729 SVs HiFi for amplified PacBio HiFi assemblies; 1017 SVs Nanopore, 888 SVs HiFi for non-amplified PacBio HiFi assemblies). All other assemblies had much fewer SVs, with second highest values for the Verkko amplified PacBio HiFi assembly (504 SVs Nanopore, 387 SVs HiFi) and the Flye non-amplified PacBio HiFi assembly (88 SVs Nanopore, 50 SVs HiFi). The addition of Nanopore reads to hifiasm assemblies of PacBio HiFi reads, amplified and non-amplified, did decrease the amounts of SVs.

### Ortholog and *k*-mer completeness support successful allele separation

We then analyzed the completeness of the assemblies based on BUSCO orthologs and *k*-mers (Figure 2). BUSCO orthologs are highly conserved genes within a target lineage, and are usually expected in single copy in a collapsed haploid assembly. For a phased assembly of a diploid species, they would instead be expected in two copies, thus duplicated. We checked both the overall BUSCO completeness (single copy and duplicated) against the Nematoda odb12 lineage, and the number of BUSCO duplicates to further evaluate phasing. As BUSCO orthologs only evaluate a restricted set of genes, we additionally investigated *k* -mer completeness, which address the whole genome content. *k* -mers are *k* nucleotide sequences extracted from a high-accuracy sequencing dataset. Based on their multiplicity, we can assess whether these *k* -mers were integrated in the contigs and in the expected number of copies. For a diploid genome, a *k* -mer spectrum should display two peaks: the first one for heterozygous *k* -mers; the second one for homozygous *k* -mers, at twice the multiplicity of the heterozygous peak, as these *k* -mers are found on the two haplotypes. Therefore, a complete diploid phased assembly should have heterozygous *k* -mers represented once, and homozygous *k* -mers represented twice. We collected the proportions of 2X homozygous *k* -mers and 1X heterozygous *k* -mers provided by KAT [43]. For the assemblies to be the most complete, both values should reach 100%. *k* -mers were identified in the Illumina sequencing dataset, as a separate sequencing technology to avoid biased assessment from comparing with the Nanopore or PacBio HiFi reads.

**Figure 2:**
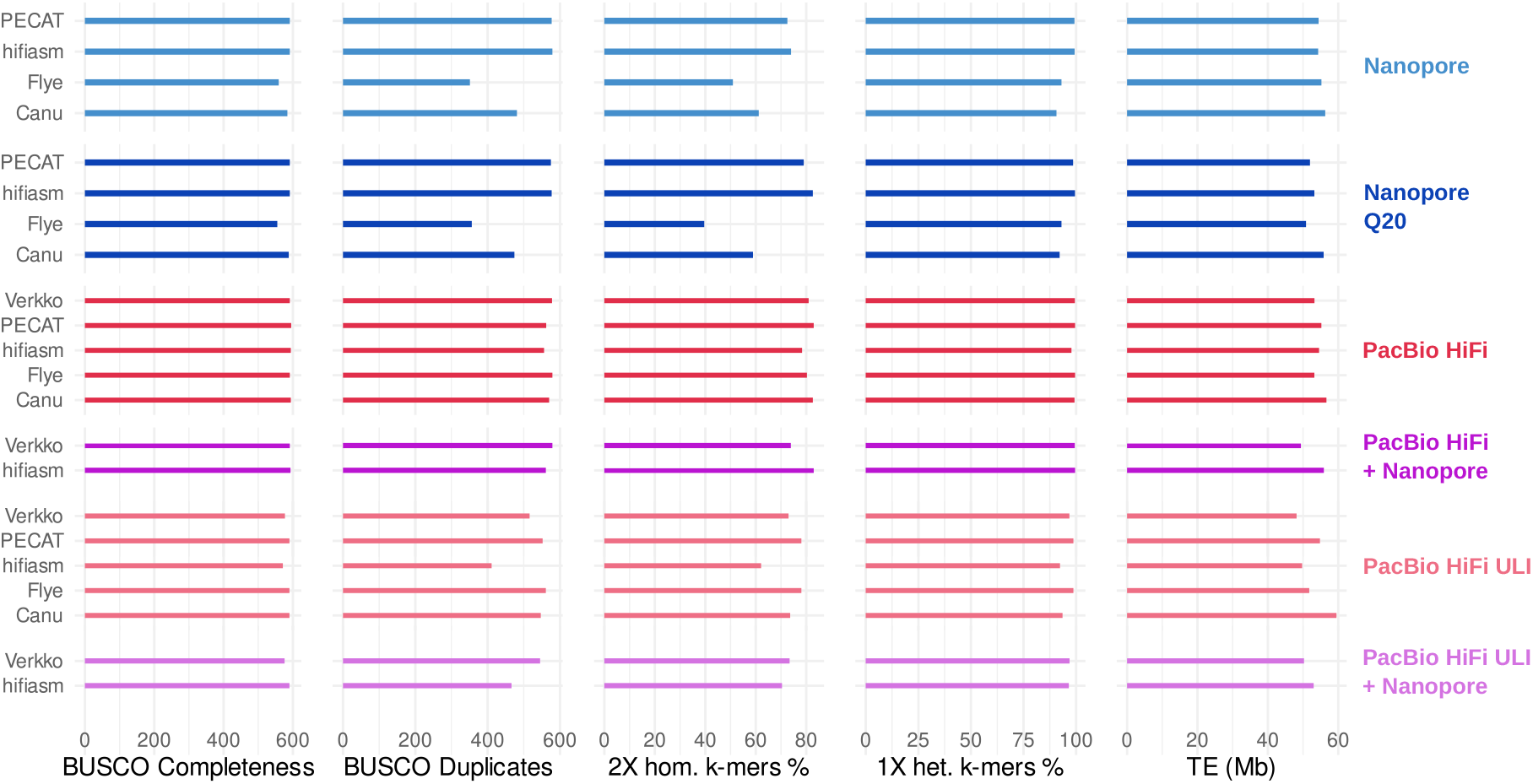
Phased assembly statistics, including the overall BUSCO completeness (single-copy and duplicated orthologs found in the assembly), the number of BUSCO duplicates (orthologs in more than one copy) against the Nematoda od12 lineage, the percentage of 2X homozygous *k* -mers and 1X heterozygous *k* -mers, and the transposable element content.

**Figure 3:**
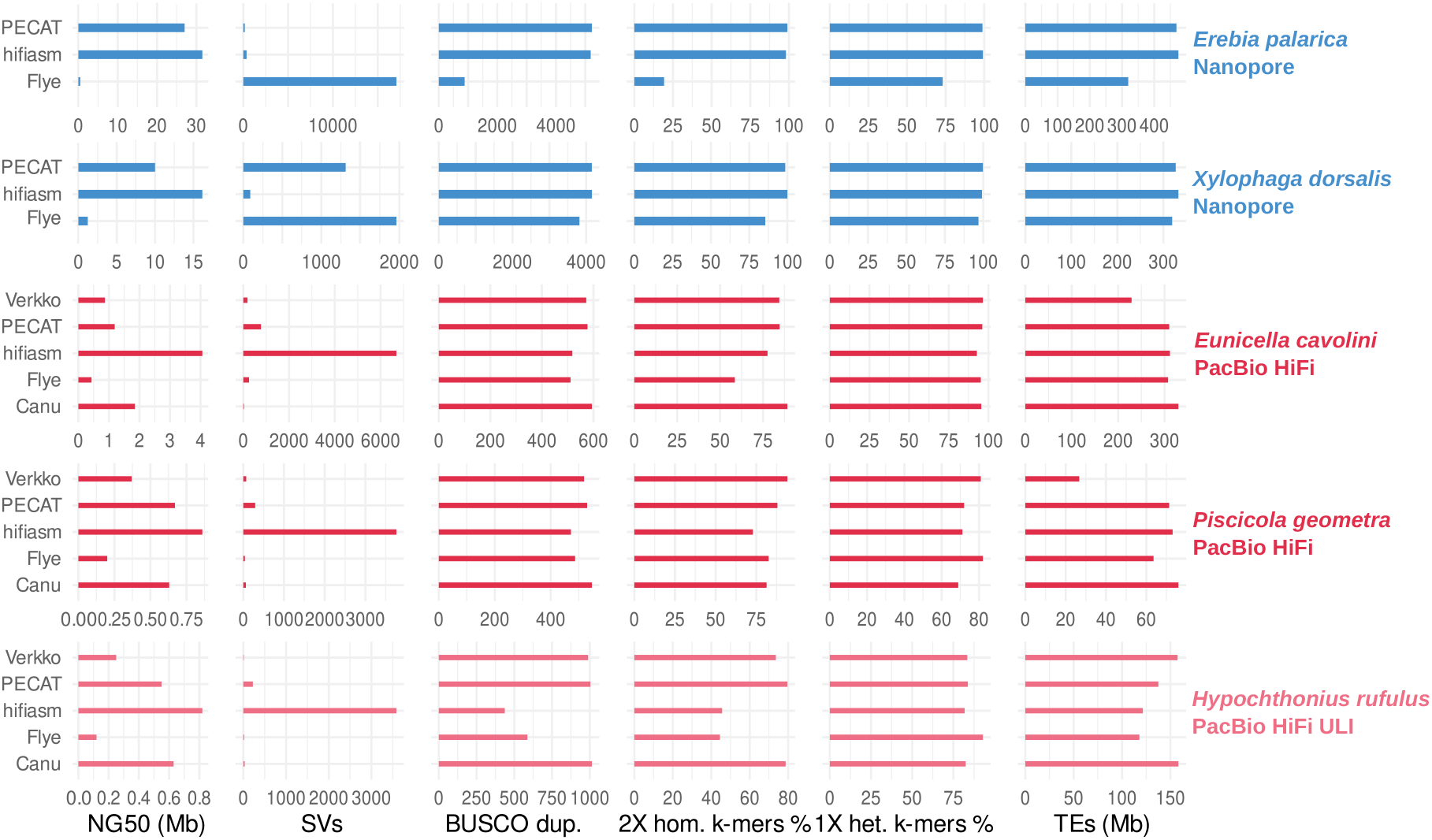
Phased assembly statistics for several non-model animal genomes, including assembly size, number of contigs, NG50, NG90, number of structural variants, number of BUSCO duplicates (orthologs in more than one copy), percentage of 2X homozygous *k* -mers and 1X heterozygous *k* -mers.

These metrics show much less variability than contiguity and SV measurements, as almost all assemblies achieved similarly high values. Out of 596 orthologs in the Nematoda odb12 lineage, a minimum of 555 and 559 orthologs overall (single copy and duplicated) was found for the Flye assemblies of the Nanopore reads and Nanopore Q20 reads. A maximum of 591 orthologs was found for hifiasm and PECAT assemblies of Nanopore reads and Nanopore Q20 reads. All non-amplified PacBio HiFi assemblies reached at least 591 orthologs, and up to 595 orthologs for the PECAT assembly. BUSCO completeness was slightly lower for amplified PacBio HiFi assemblies, from 571 orthologs for hifiasm to 590 orthologs for Canu, Flye and PECAT assemblies.

BUSCO duplicates highlight less properly phased assemblies (Figure 2). Flye and Canu assemblies under-performed for Nanopore and Nanopore Q20 assemblies with numbers of duplicates between 351 (Flye) and 481 (Canu), compared to hifiasm and PECAT assemblies at 575-579 duplicated orthologs, which represents almost all identified orthologs. Non-amplified PacBio HiFi assemblies all had over 556 duplicated orthologs, with a lowest value for the hifiasm assembly, and the highest value for Flye at 579 orthologs, followed by Verkko with 578 duplicated orthologs (and 579 when combined with Nanopore reads). Amplified PacBio HiFi assemblies had lower numbers of duplicates, with a minimum value of 411 for hifiasm, and a maximum of 561 for Flye.

*k* -mer completeness mirrors ortholog completeness (Figure 2, Supplementary Figures S3-8). Flye and Canu assemblies of Nanopore reads and Nanopore Q20 reads have smaller proportions of 2X homozygous *k* -mers and 1X heterozygous *k* -mers than other assemblers. Proportions of 2X homozygous *k* -mers in Nanopore assemblies are overall lower than for non-amplified PacBio HiFi assemblies, with a maximum of 73.95% for hifiasm (Nanopore), versus 78.33 to 82.96% for hifiasm, and PECAT assemblies (non-amplified PacBio HiFi). However, this gap narrows when using only Nanopore Q20 reads, as hifiasm Nanopore Q20 reached 82.60% and PECAT Nanopore Q20 79.01%. With amplified PacBio HiFi, the hifiasm assembly had the lowest proportion of 2X homozygous *k* -mers at 62.11%, while Flye and PECAT culminated at 78.06%. Regarding 1X heterozygous *k* -mers, all values were above 90.66% (Canu Nanopore). The highest percentages for Nanopore assemblies were found for hifiasm and PECAT, reaching over 98.55% with Nanopore reads and Nanopore Q20 reads. All non-amplified PacBio HiFi assemblies reached a minimum of 97.78% (hifiasm); amplified PacBio HiFi resulted in slightly smaller percentages, down to 92.34% for hifiasm, but with a maximum of 98.72% for Flye and PECAT.

### Similar transposable element content between sequencing technologies and assemblers

To further evaluate assembly completeness and the representation of ‘genomic dark matter,’ we characterized the repetitive content, specifically focusing on transposable elements (TEs). The structural resolution of TEs has been significantly enhanced by long-read technologies, which overcome the inherent limitations of short reads in spanning complex, repetitive segments [29]. As the groundtruth amount of TEs in *Plectus sambesii* is not known, we focused on identifying consistent trends across assembly strategies. Furthermore, we analyzed total TE content in absolute base pairs rather than as a percentage of the assembly; this avoids the biases introduced by variable assembly sizes and ensures a more direct comparison of the repetitive sequence recovered by each tool. The median amount of repetitions across sequencing technologies and assemblers reaches 53.8 Mb. Assemblies of either Nanopore reads or non-amplified PacBio HiFi reads have similar TE contents ranging from 53.1 Mb and 56.6 Mb. When using Nanopore Q20 reads only, Flye and PECAT yielded lower amounts of TEs (50.8 and 52.0 Mb). TE content was the most variable in assemblies of amplified PacBio HiFi reads, from 48.2 (Verkko) to 59.5 Mb (Canu). This may be explained by amplification bias leading to variations in coverage that impede the representation of repetitive sequences in their correct number. The assembler Canu yielded the highest amount of transposable elements across all long-read technologies, with increased values in each category. This suggests that Canu was either more efficient at resolving TEs than other assemblers, or tends to overrepresent them. Verkko led to smaller TE content, and in addition, higher amounts of non-TIR elements, repeat regions, coupled with lower amounts of TIR elements (Supplementary Table S1). This could suggest difficulties for EDTA to classify TIR, non-TIR elements and repeats regions in the way that they were included in the Verkko assemblies, or poor resolution of these categories of transposable elements.

### Consistent assembly results across diverse animal species

Assembly evaluation for *Plectus sambesii* highlighted several strengths and flaws depending on sequencing technologies and assembler. To investigate whether these results hold beyond the specific genome of *Plectus sambesii*, we tested assemblers on multiple non-model diploid animal species for which one long-read sequencing dataset was available: Nanopore R10.4.1 reads for the butterfly *Erebia palarica* [48] and the bivalve *Xylophaga dorsalis* [35]; non-amplified PacBio HiFi reads for the Gorgonian *Eunicella cavolini* [49] and the fish leech *Piscicola geometra* [36]; amplified PacBio HiFi reads for the mite *Hypochthonius rufulus* [37] (Table 1). The first four datasets were generated as part of the Darwin Tree of Life [14] or the European Reference Genome Atlas [12]. They represent diverse branches of metazoans, with different coverage depth, heterozygosity, and repeat content. We investigated assembly size, contiguity, structural correctness, ortholog and *k* -mer completeness, and transposable element content.

**Table 1:** Genomic features of non-model animal species included in this study. The depth refers to the multiplicity of the heterozygous peak identified by KAT, and represents the coverage depth for each haplotype. The estimated genome size is based on the published collapsed assemblies; we expect double this size for a phased assembly of a diploid species. The heterozygosity was computed using GenomeScope2 (Supplementary Figure S9)). Repeat content is based on published assemblies and EDTA annotation.

| Species | Sequencing data | Depth | Estimated size (Mb) | Het. % | Repeat % |
| --- | --- | --- | --- | --- | --- |
| *Plectus sambesii* | Nanopore R10.4.1 | 46 | 120 | 3.8 | 22.4 |
|  | PacBio HiFi | 27 |  |  |  |
|  | PacBio HiFi ULI | 40 |  |  |  |
| *Erebia palarica* | Nanopore R10.4.1 | 84 | 513 | 1.0 | 49.6 |
| *Xylophaga dorsalis* | Nanopore R10.4.1 | 83 | 452 | 3.1 | 44.2 |
| *Eunicella cavolini* | PacBio HiFi | 24 | 386 | 2.2 | 36.8 |
| *Piscicola geometra* | PacBio HiFi | 37 | 205 | 1.1 | 16.1 |
| *Hypochthonius rufulus* | PacBio HiFi ULI | 101 | 185 | 1.2 | 41.1 |

Patterns observed in *Plectus sambesii* are found similarly in these multiple species, with regards for sequencing technology and assembler. Assemblies of Nanopore reads with hifiasm and PECAT for *Erebia palarica* and *Xylophaga dorsalis* yielded larger assembly sizes than with Flye (959-961 Mb versus 637 Mb, Supplementary Tables S2-3). Considering that the expected haploid genome size was estimated to 401 Mb (Supplementary Figure S9) and the assembly size of 513 Mb of the reference collapsed assembly [48], the Flye assembly is far shorter than the expected diploid size. hifiasm and PECAT assemblies also had higher rates of duplicated BUSCO orthologs (5,184-5,226 over 5,760 orthologs), 1X heterozygous (99.12-98.81%) and 2X homozygous (98.06-99.01%) *k* -mers, indicating that hifiasm and PECAT separated haplotypes well (Supplementary Figures S10-11). hifiasm and PECAT assemblies reached proportions of both 1X heterozygous and 2X homozygous *k* -mers beyond 98%, indicating a high completeness of the two haplotypes. Along high ortholog and *k* -mer completeness, hifiasm and PECAT also yielded more repetitive content than Flye. hifiasm had overall the highest amount of TEs for both *E. palarica* (475.0 Mb) and *X. dorsalis* (332.7 Mb), however, this is only because of the high amount of LTRs (Supplementary Tables S2-3). For all other categories, the largest amount of TEs was identified in PECAT assemblies. hifiasm resulted in the lowest amount of structural variants (361 SVs for *E. palarica*, 89 SVs for *dorsalis*), based on Nanopore reads, far below the values of Flye and PECAT (17,080 and 1,312 SVs for *E. palarica*, 1,962 and 1,312 SVs for *X. dorsalis*).

Assemblies of non-amplified PacBio HiFi reads for *Eunicella cavolini* had similar sizes (802 to 862 Mb). hifiasm achieved the highest NG50 (4.1 Mb), followed by Canu (1.9 Mb) and PECAT (1.2 Mb). The Canu assembly also had the highest number of BUSCO duplicates (594 out of 672 orthologs) and proportion of 2X homozygous *k* -mers (89.22%) (Supplementary Figure S12). Except for hifiasm at 92.77%, all assemblers yielded contigs with a proportion of 1X heterozygous *k* -mers over 95%. Low number of BUSCO duplicates were observed for the Flye and hifiasm assemblies (511 and 518 orthologs), along lower proportions of 2X homozygous *k* -mers (58.60 and 77.62%). A lower amount of TE was found in the Flye assembly of *P. geometra*, at 63.6 Mb. For *Piscicola geometra*, results were again similar to those observed for non-amplified PacBio HiFi assemblies of *Plectus sambesii* and *Eunicella cavolini*. The highest NG50 was obtained again using hifiasm (0.9 Mb), followed by PECAT (O.7 Mb) and Canu (0.6 Mb). Still, the hifiasm assembly was less complete than other assemblies, with 472 duplicated orthologs (out of 672 orthologs), 71.12% of 1X heterozygous *k* -mers and 72.88% of 2X homozygous *k* -mers (Supplementary Figure S13). PECAT and Canu yielded the largest number of duplicated BUSCO orthologs (530 and 547), and Flye and Verkko performed the best regarding *k* -mer completeness. The highest quantity of Tes was found in Canu assemblies, for all categories, and much lower amounts were found in Verkko assemblies as observed in *Plectus sambesii* (Supplementary Tables S4-5). Comparable TE contents were found in hifiasm and PECAT assemblies (311.5 and 310.0 Mb for *E. cavolini*, 73.1 and 71.3 Mb for *P. geometra*). The highest amount of SVs was found for hifiasm (6,690 SVs for *E. cavolini*, 3,762 SVs for *P. geometra*), and the second highest for PECAT (773 for *E. cavolini*, 292 SVs for *P. geometra*).

Assemblies of amplified PacBio HiFi reads of *Hypochthonius rufulus* led to high numbers of BUSCO duplicates (989 to 1014 out of 1123 orthologs), along high proportions of 1X heterozygous and 2X homozygous *k* -mers (Supplementary Figure S14). However, low completeness was observed in Flye and hifiasm assemblies, regarding BUSCO duplicates (587 and 437) and 2X homozygous *k* -mers (45.62 and 44.50%). TE content ranged from 118.0 Mb (Flye) to 158.3 Mb (Canu), with a second highest value for Verkko at 157.4 Mb (Supplementary Table S6). 3,584 SVs were found for the hifiasm assembly, compared to 224 SVs for PECAT and below for other assemblers. For all assemblies and across all five species, as observed in *Plectus sambesii*, SVs consisted mostly of insertions and deletions (Supplementary Tables S2-6).

## Discussion

Collapsed assemblies were a standard for reference assemblies based on low-accuracy long reads. High-accuracy long reads are taking genome assemblies one step further by bringing a new era of phased assemblies, paving the way for innovative analyses addressing haplotype coexistence. Our results show that haplotype-resolved assemblies are now achievable for a variety of non-model species, and provide a more comprehensive representation of genomes with mid to high heterozygosity.

To compare genome assemblies of these diverse animal species, we conducted a thorough evaluation by repurposing state-of-the-art programs to collect haplotype-relevant statistics, going beyond the initial assessment of contiguity. We assessed phasing efficiency by examinating structural correctness through SV calling, ortholog and *k* -mer completeness, and transposable element content. Contiguity, orthologs, and *k* -mer representation are already commonly included in evaluation steps. By incorporating SV detection in our analysis, we observed that high contiguity does not necessarily correlate with structural accuracy. Notably, some assemblies exhibiting superior NG50 also displayed a disproportionately high number of indels, whereas other assemblers produced shorter but more accurate contigs. In addition, the analysis of transposable element content revealed that assembly strategy can affect the representation of the repetitive landscape, thereby impacting downstream interpretations. Consequently, the evaluation of SV profiles, alongside repetitive content recovery, provides a crucial diagnostic layer to distinguish truly robust assemblies from those where contiguity is achieved at the expense of base-level or structural correctness.

We observed robust phasing efficiency among PacBio HiFi assemblers. Although hifiasm is often preferred in PacBio HiFi assembly pipelines and consistently yielded the highest NG50s, Canu and PECAT reached higher ortholog and *k* -mer completeness, with fewer structural inaccuracies. While assemblies of non-amplified PacBio HiFi reads neared chromosome-level contigs, amplified PacBio HiFi reads led to a tradeoff in assembly contiguity. However, their NG50s over 1 Mb were far over the contiguities which could be achieved in short-read assemblies. Furthermore, the completeness and repetitive content of amplified PacBio HiFi assemblies reached values close to those of assemblies from non-amplified reads. The generation of high-quality assemblies from less than a nanogram of DNA constitutes a major technological leap for the study of non-model species. Indeed, the amount of DNA required as input is often a limiting factor for species with small individuals; thus the possibility to produce assemblies from a single individual may be what allows a research question to be explored or not.

A key finding of this study is the remarkable performance of Nanopore R10.4.1 data, which, when paired with assemblers like hifiasm or PECAT, yield results comparable to the established PacBio HiFi gold standard. Both hifiasm and PECAT assemblies of Nanopore reads reached high contiguity, completeness, and structural correctness. The outstanding haplotype resolution of hifiasm assemblies shows that the innovative read-phasing correction approach implemented in the Nanopore-adapted version is particularly well designed for haplotype separation.

Crucially, our results reveal that the choice of sequencing technology is no longer the primary bottleneck for phasing feasibility. Nanopore R10.4.1 data demonstrated that they can match the structural fidelity of PacBio HiFi when paired with the most appropriate assembler. This is a major finding for the global democratization of genomics, to enable sequencing and assembly of high-quality, phased references both in large hubs and in smaller, local infrastructures.

Based on our benchmarking of diverse non-model animal species, we provide a clear decision framework, and propose a novel approach to genome assembly projects, prioritizing a phased-first paradigm. This will not only ensure the biological integrity of the data used in biodiversity genomics but also empower a truly global community to document the complexity of life with unprecedented precision.

## Data & Code availability

Datasets were deposited on the European Nucleotide Archive under the Bioproject PRJEB74285. More precisely, the reads are available under run accession numbers: Nanopore R10.4.1 ERR12861248; non-amplified PacBio HiFi ERR17019299, ERR17019300, ERR17019301, ERR17019600; single-individual amplified PacBio HiFi ERR17017482. Command lines are provided at github.com/nadegeguiglielmoni/phasing_assemblies.

## Acknowledgements

We thank Laura I. Villegas for her contribution to long-read sequencing. We additionally thank Anja Schuster, Kerstin Becker and the Genomics and Transcriptomics Laboratory of the Heinrich Heine University Düsseldorf for the generation of PacBio HiFi sequencing data.

