## Supplementary_Figures for "Phasing genome assemblies of non-model animal species in the era of high-accuracy long reads"


(a) GenomeScope (b) Nanopore

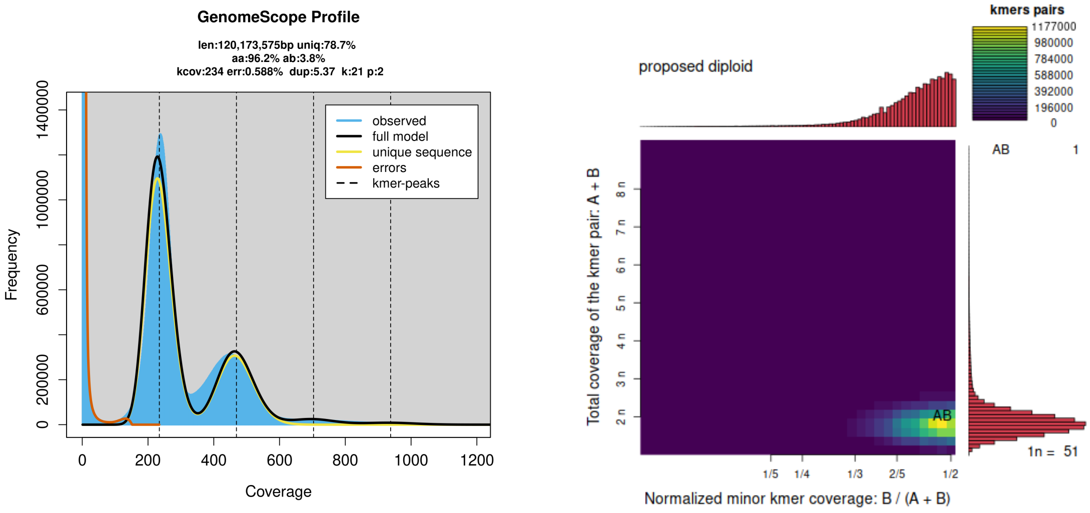


Figure 1: $k$-mer analysis of Illumina reads.

(a) Nanopore (b) PacBio HiFi (c) ULI PacBio HiFi

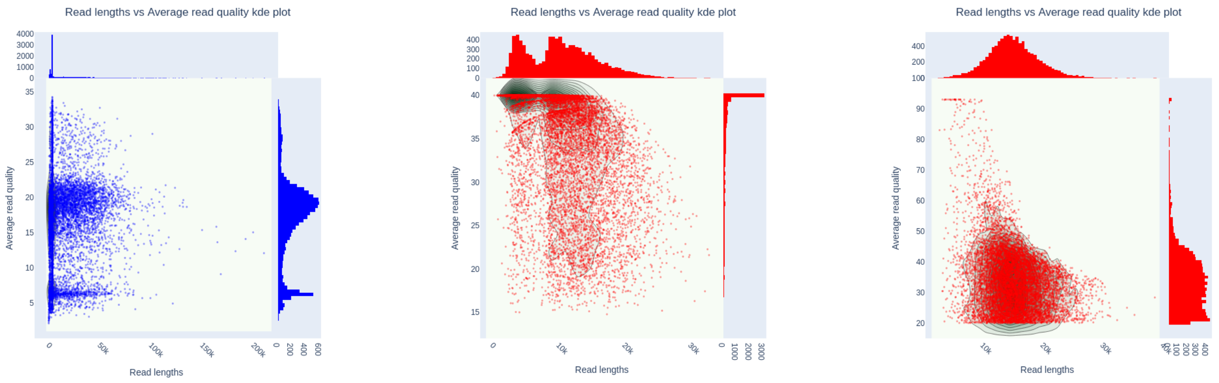


Figure 2: NanoPlot analysis of the Nanopore, non-amplified PacBio HiFi and ULI PacBio HiFi reads.

(a) Canu (b) Flye

(c) hifiasm (d) PECAT

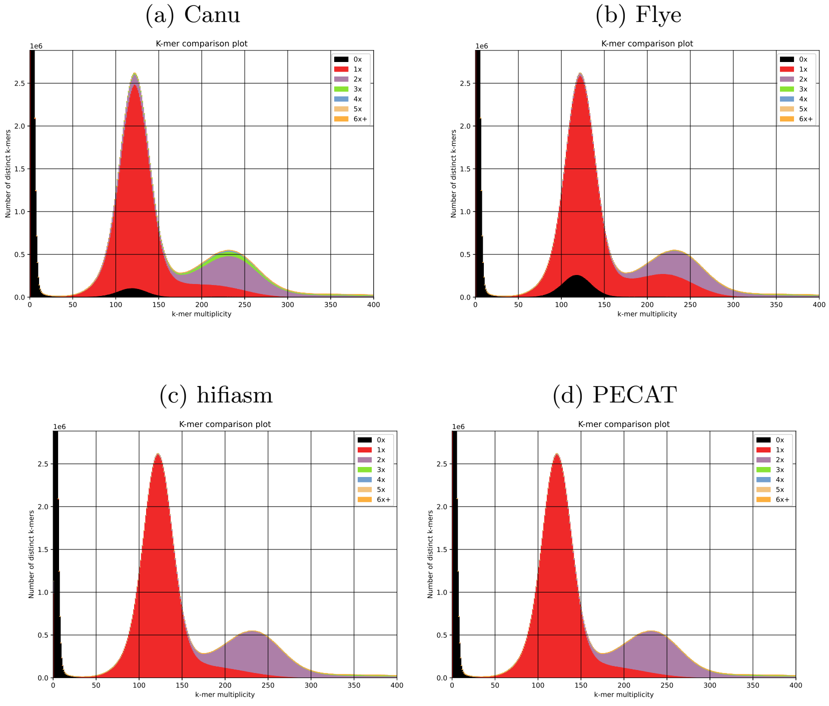


Figure 3: KAT analysis of Nanopore assemblies for *Plectus sambesii*.

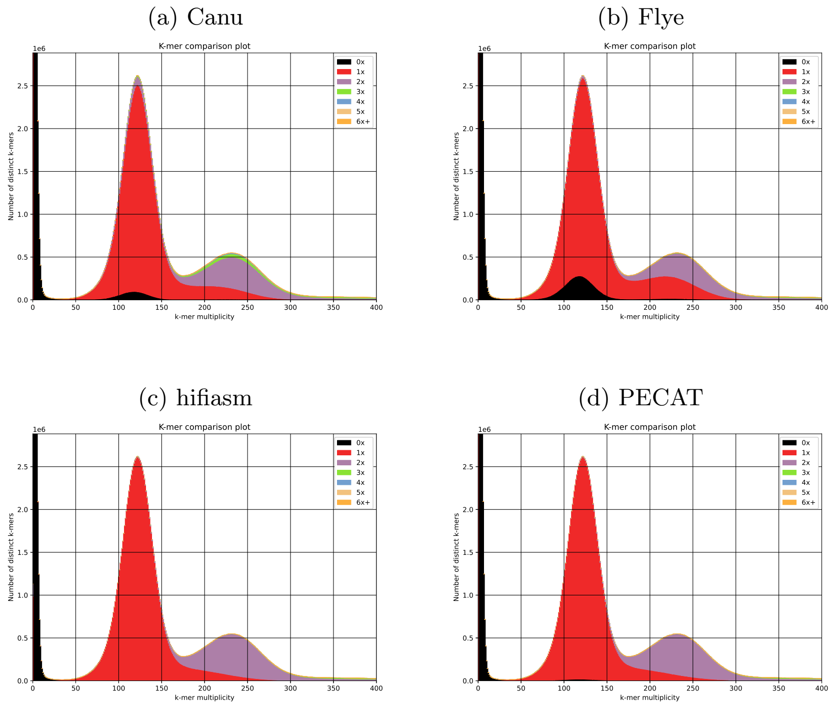


Figure 4: KAT analysis of Nanopore Q20+ assemblies for *Plectus sambesii*.

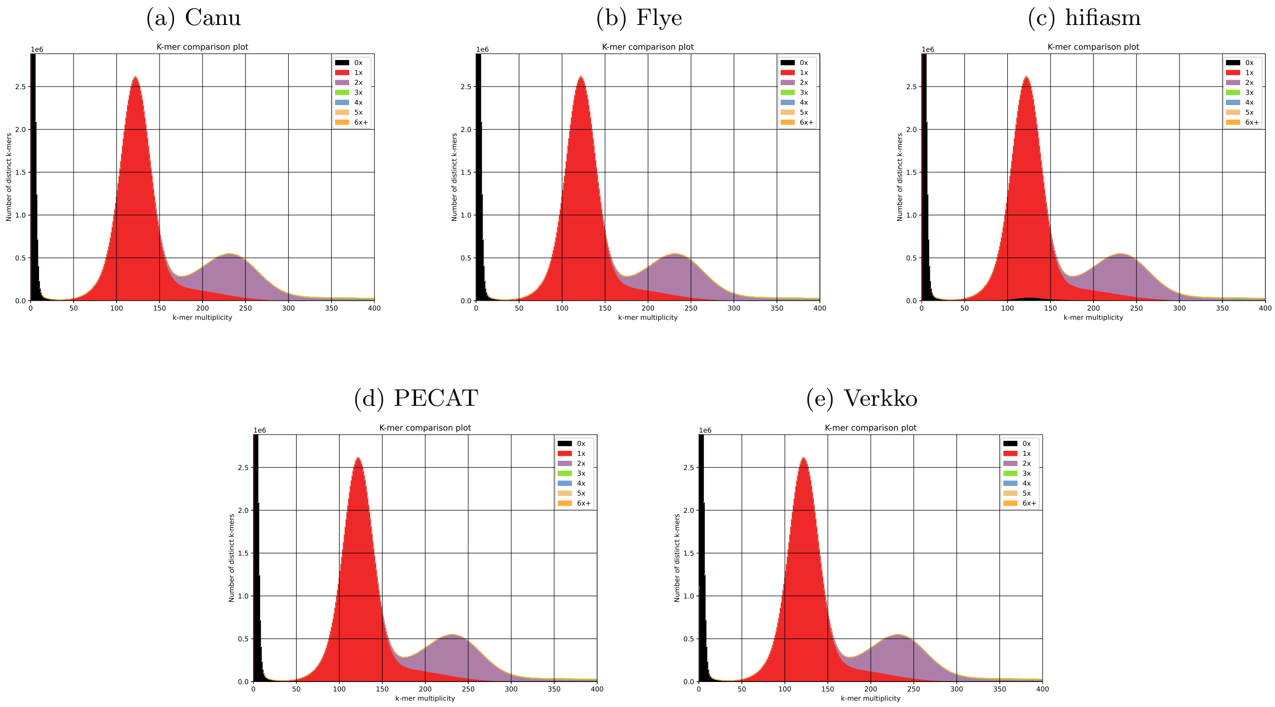


Figure 5: KAT analysis of PacBio HiFi assemblies for *Plectus sambesii*.

(a) hifiasm

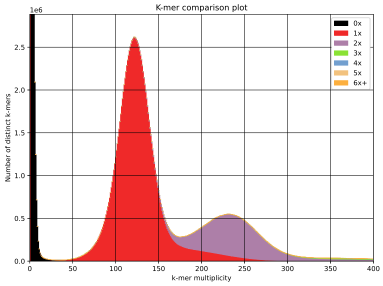


(b) Verkko

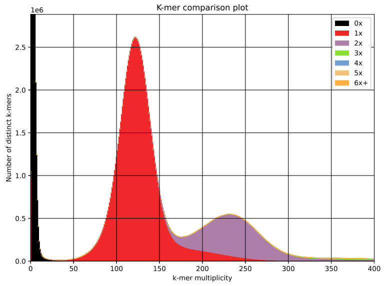


Figure 6: KAT analysis of PacBio HiFi + Nanopore assemblies for *Plectus sambesii*.

(a) Canu

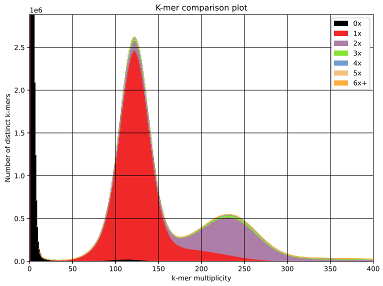


(b) Flye

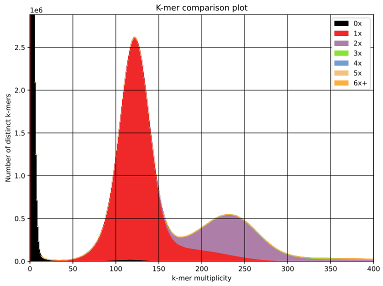


(c) hifiasm

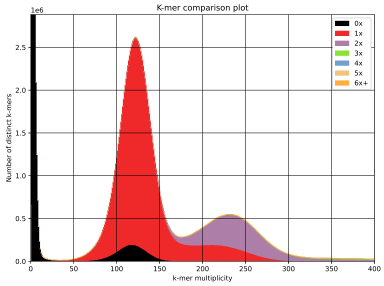


(d) PECAT

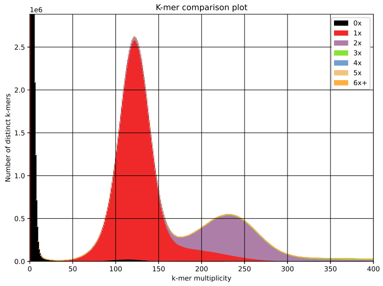


(e) Verkko

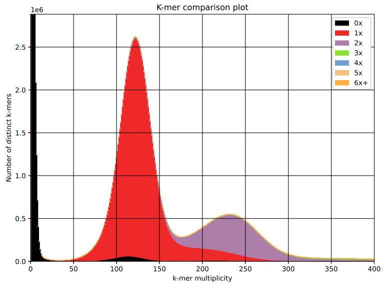


Figure 7: KAT analysis of ULI PacBio HiFi assemblies for *Plectus sambesii*.

(a) hifiasm

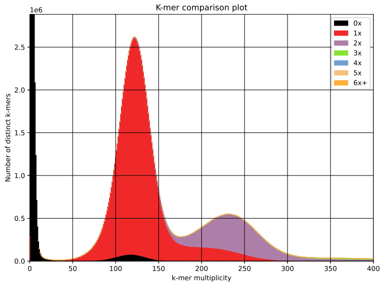


(b) Verkko

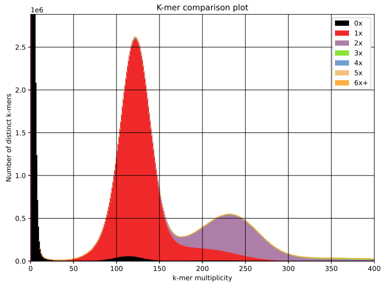


Figure 8: KAT analysis of ULI PacBio HiFi + Nanopore assemblies for *Plectus sambesii*.

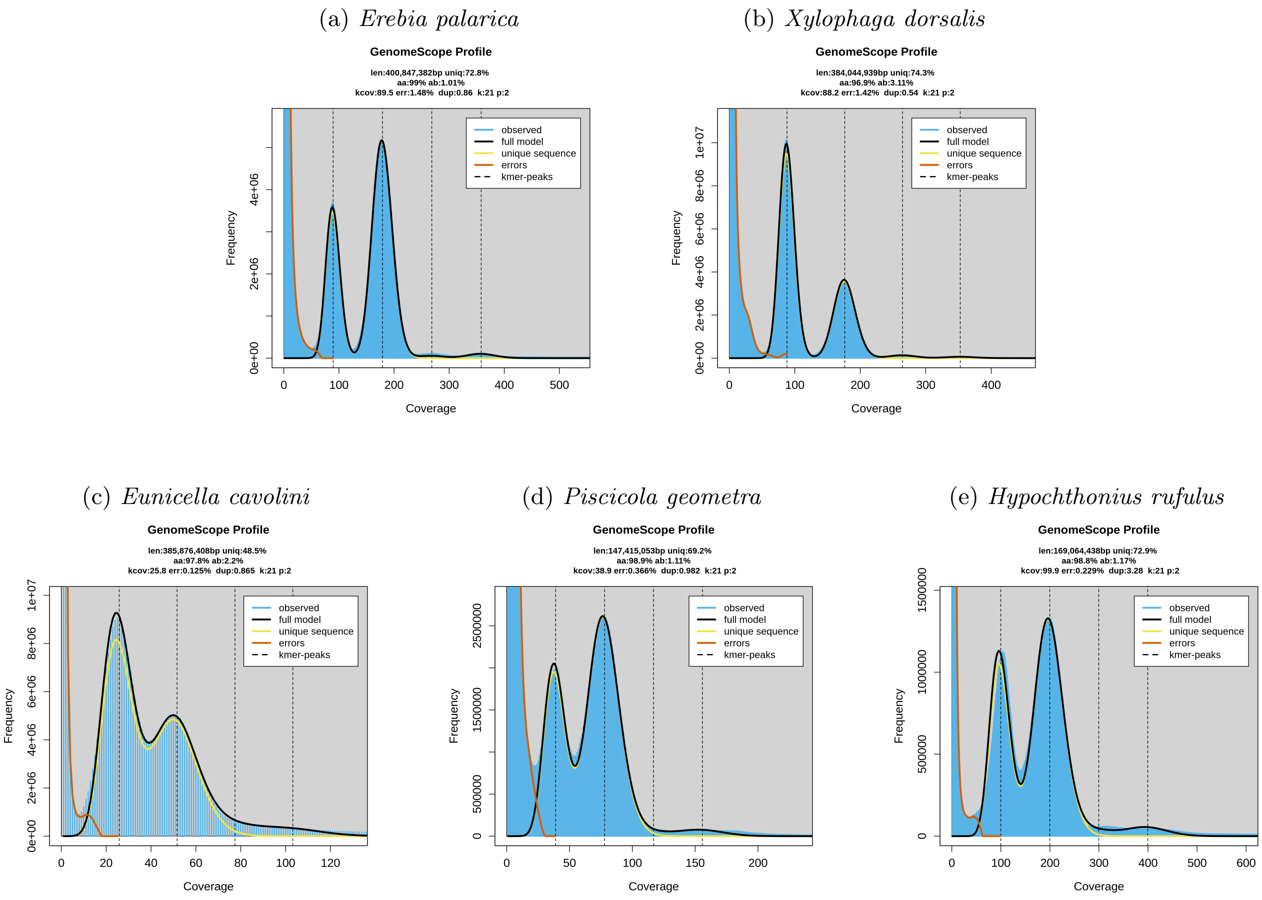


Figure 9: GenomeScope2 analysis of *Erebia palarica* and *Xylophaga dorsalis* based on Nanopore reads, *Eunicella cavolini* and *Piscicola geometra* based on PacBio HiFi reads, and *Hypochthonius rufulus* based on ULI PacBio HiFi reads.

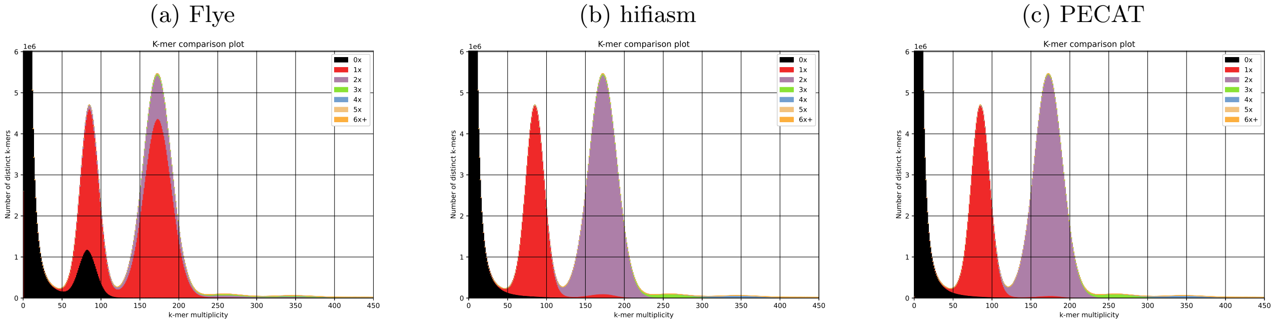


Figure 10: KAT analysis of Nanopore assemblies for *Erebia palarica*.

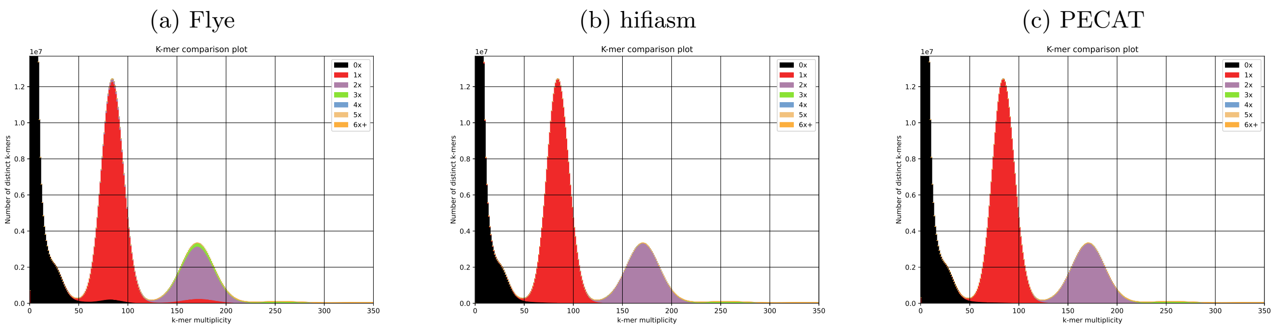


Figure 11: KAT analysis of Nanopore assemblies for *Xylophaga dorsalis*.

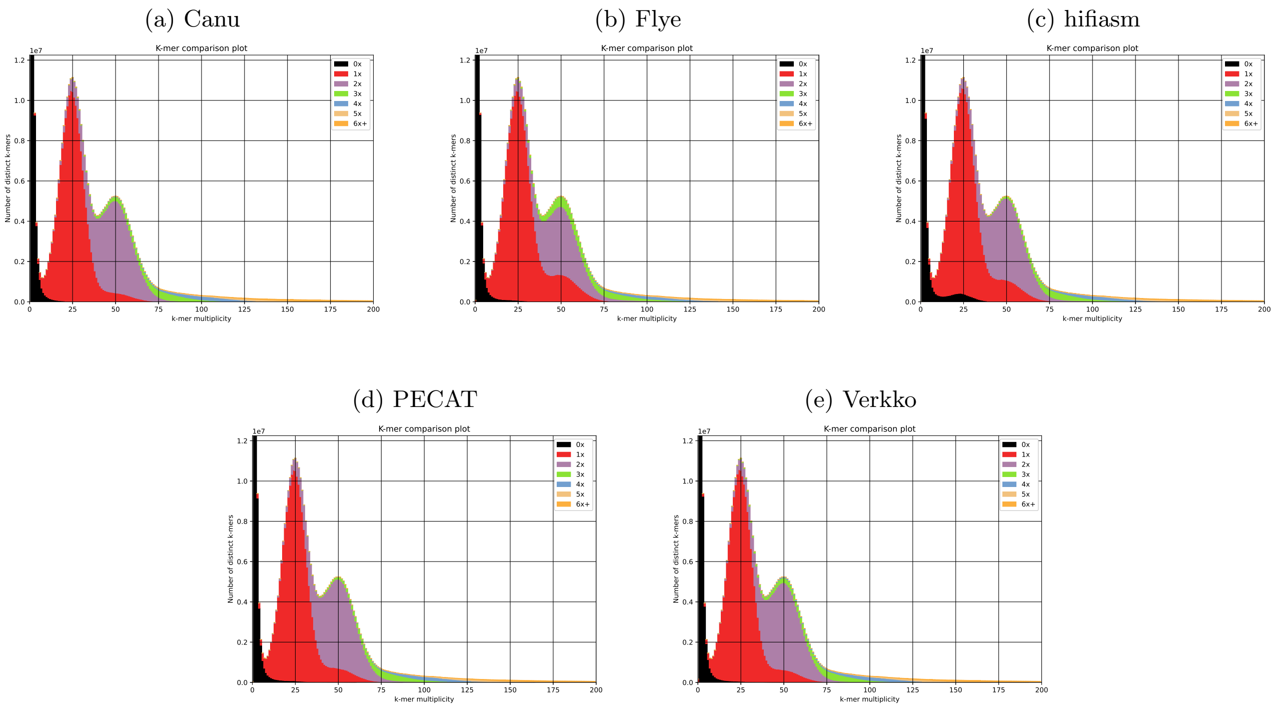


Figure 12: KAT analysis of PacBio HiFi assemblies for *Eunicella cavolini*.

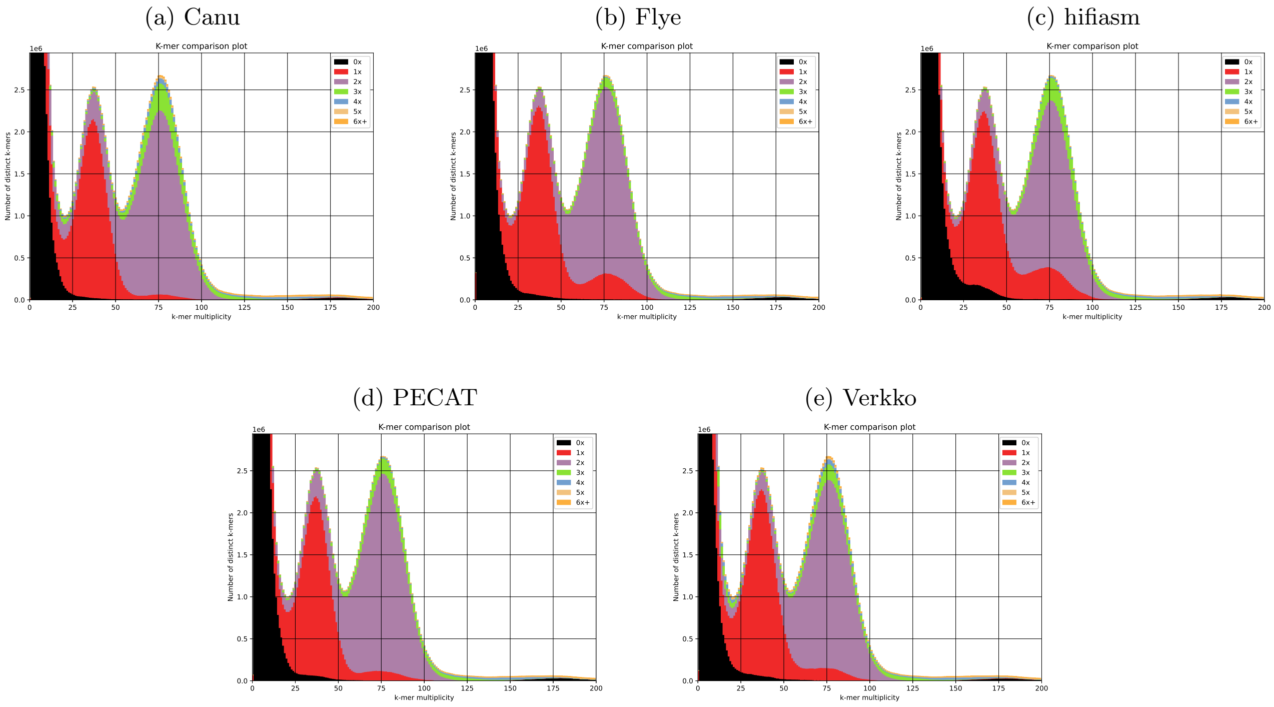


Figure 13: KAT analysis of PacBio HiFi assemblies for *Piscicola geometra*.

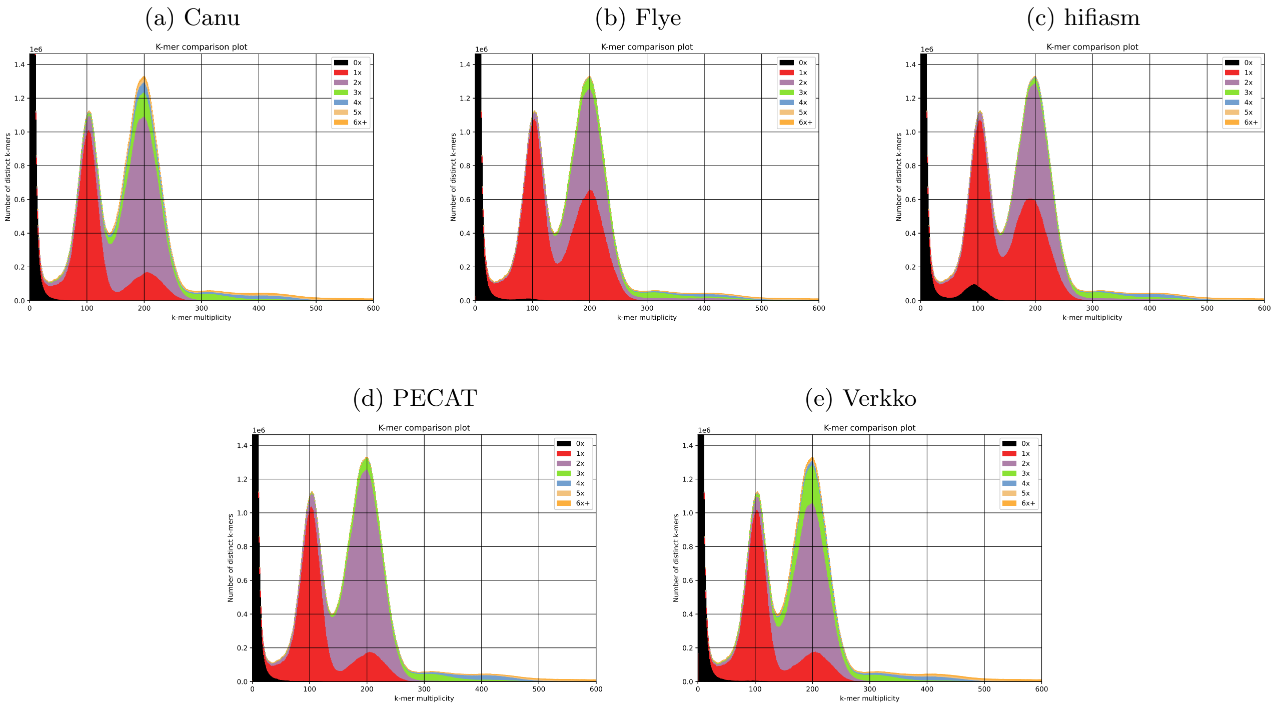


Figure 14: KAT analysis of PacBio HiFi assemblies for *Hypochthonius rufulus*.
