## Supplementary_Tables for "Phasing genome assemblies of non-model animal species in the era of high-accuracy long reads"

Table 1: Detailed assembly statistics for Nanopore, Nanopore Q20, non-amplified PacBio HiFi, and ULI PacBio HiFi assemblies of *Plectus sambesii*. These statistics include the assembly size, contig number, NG50, NG90, number of single and duplicate BUSCO orthologs, proportions of 1X heterozygous *k*-mers and 2X homozygous *k*-mers, amount of transposable elements per category, and amount of structural variants per category identified with Nanopore and non-amplified PacBio HiFi reads.

Table 2: Detailed assembly statistics for Nanopore assemblies of *Erebia palarica*. These statistics include the assembly size, contig number, NG50, NG90, number of single and duplicate BUSCO orthologs, proportions of 1X heterozygous $k$-mers and 2X homozygous $k$-mers, amount of transposable elements per category, and amount of structural variants per category identified with Nanopore reads.

Table 3: Detailed assembly statistics for Nanopore assemblies of *Xylophaga dorsalis*. These statistics include the assembly size, contig number, NG50, NG90, number of single and duplicate BUSCO orthologs, proportions of 1X heterozygous $k$-mers and 2X homozygous $k$-mers, amount of transposable elements per category, and amount of structural variants per category identified with Nanopore reads.

Table 4: Detailed assembly statistics for PacBio HiFi assemblies of *Eunicella cavolini*. These statistics include the assembly size, contig number, NG50, NG90, number of single and duplicate BUSCO orthologs, proportions of 1X heterozygous $k$-mers and 2X homozygous $k$-mers, amount of transposable elements per category, and amount of structural variants per category identified with PacBio HiFi reads.

Table 5: Detailed assembly statistics for PacBio HiFi assemblies of *Piscicola geometra*. These statistics include the assembly size, contig number, NG50, NG90, number of single and duplicate BUSCO orthologs, proportions of 1X heterozygous $k$-mers and 2X homozygous $k$-mers, amount of transposable elements per category, and amount of structural variants per category identified with PacBio HiFi reads.

Table 6: Detailed assembly statistics for ULI PacBio HiFi assemblies of *Hypochthonius rufulus*. These statistics include the assembly size, contig number, NG50, NG90, number of single and duplicate BUSCO orthologs, proportions of 1X heterozygous $k$-mers and 2X homozygous $k$-mers, amount of transposable elements per category, and amount of structural variants per category identified with ULI PacBio HiFi reads.
